# Immune Inhibitor A Metalloproteases Contribute to Virulence in *Bacillus* Endophthalmitis

**DOI:** 10.1101/2020.07.06.190264

**Authors:** Erin T. Livingston, Md Huzzatul Mursalin, Phillip S. Coburn, Roger Astley, Frederick C. Miller, Omar Amayem, Didier Lereclus, Michelle C. Callegan

## Abstract

Bacterial endophthalmitis is a devastating infection that can cause blindness following the introduction of organisms into the posterior segment of the eye. Over half of *Bacillus* endophthalmitis cases result in significant loss of useful vision. Often, these eyes have to be enucleated. *Bacillus* produces many virulence factors in the eye that may contribute to retinal damage and robust inflammation. This study analyzed *Bacillus* immune inhibitor A (InhA) metalloproteases, which digest extracellular matrix, tight junction proteins, and antimicrobial proteins. We hypothesized that InhAs contribute to *Bacillus* intraocular virulence and inflammation. We analyzed phenotypes and infectivity of wild type (WT), InhA1-deficient (Δ*inhA1*), InhA2-deficient (Δ*inhA2*), or InhA1, A2, and A3-deficient (Δ*inhA1-3*) Bacillus thuringiensis. *In vitro* analysis of growth, proteolysis, and cytotoxicity were compared between *B. thuringiensis* strains. WT and InhA mutants were similarly cytotoxic to retinal cells. Mutant Δ*inhA1* and Δ*inhA2* entered log phase growth earlier than WT. Proteolysis of the Δ*inhA1-3* mutant was decreased, but this strain grew similar to WT *in vitro*. Experimental endophthalmitis was initiated by intravitreally infecting C57BL/6J mice with 200 CFU of *B. thuringiensis* WT or InhA mutants. Intraocular *Bacillus* and retinal function loss were quantified. Intraocular myeloperoxidase concentrations were quantified and histology was analyzed. Eyes infected with Δ*inhA1* or Δ*inhA2* strains contained greater numbers of bacteria than eyes infected with WT throughout the course of infection. Eyes infected with single mutants had inflammation and retinal function loss similar to eyes infected with WT strain. Eyes infected with Δ*inhA1-3* cleared the infection, with less retinal function loss and inflammation compared to eyes infected with the WT strain. RT-PCR results suggested that single InhA mutant results may be explained by compensatory expression of the other InhAs in these mutants. These results indicate that together, the InhA metalloproteases contribute to the severity of infection and inflammation in *Bacillus* endophthalmitis.

**Author summary:** Bacterial endophthalmitis is an infection of the eye, which can follow accidental contamination of the posterior segment following ocular surgery (postoperative), a penetrating wound (post-traumatic), or during spread of bacteria into the eye from the bloodstream (endogenous). During bacterial endophthalmitis, virulent pathogens such as *Bacillus* cause ocular damage via the activities of an array of virulence factors, including proteases. A class of proteases that are expressed by *Bacillus* during ocular infection are the immune inhibitor A metalloproteases. Here, we used a mouse model of endophthalmitis to test mutant *Bacillus* that lack single or multiple InhAs to determine if these metalloproteases contributed to the virulence during the disease. In the absence of the production of all InhAs, *Bacillus* could not cause severe infection. Our study provides new insights into the virulence of *Bacillus* in the eye, and the contribution of its InhA metalloproteases to establishing infection.

## Introduction

One of the most severe forms of intraocular inflammation and rapid vision loss caused by bacteria is due to infection with *Bacillus* spp. [1–6]. *Bacillus* endophthalmitis occurs most often following ocular trauma involving a foreign body contaminated with this bacterium [7–11]. Despite treatment with antibiotics, anti-inflammatory drugs, and surgical intervention, more than 70% of patients with *Bacillus* endophthalmitis have been documented to have significant vision loss, and about 50% of those patients underwent evisceration or enucleation of the infected eye [7–11]. Because this feared infection is difficult to treat, there is great importance in identifying virulence factors of *Bacillus* that contribute to this blinding disease.

*Bacillus thuringiensis* belongs to the *Bacillus cereus sensu lato* group, and is known for causing severe bacterial endophthalmitis [1,12]. *B. thuringiensis* is so genetically similar to *B. cereus* that the species delineation between the two within the *sensu lato* group has been problematic despite the various approaches and techniques used [13]. Studies comparing the genomes of both organisms have suggested that they belong to the same species [14]. It has also been reported that the genetic and phenotypic properties between these bacteria are barely distinguishable [15]. Both organisms replicate quickly in the eye, are highly motile, and express similar virulence factors—all of which may contribute to the severity of endophthalmitis [12,16–19].

The majority of extracellular *Bacillus* virulence factors are produced under the control of a global regulator, PlcR. An absence of a functioning PlcR system delayed the damage typically seen in *Bacillus* endophthalmitis [16,19–21]. In intraocular infections with mutants lacking a functional PlcR system, there was still retinal toxicity, function loss, and vascular permeability. Virulence factors outside of *plcR* regulation that may have contributed to this delayed response might include cell wall endopeptidases, S-layer, hemolysins, InhA1 metalloprotease, amidases, pili, and/or flagella components [20–27]. Our previous work showed that individual toxins, such as hemolysin BL, phosphatidycholine-specific phospholipase C (PC-PLC), or phosphatidylinositol-specific phospholipase C (PI-PLC), contributed little to the disease [17,18]. Recently, we observed that specific virulence factors are highly expressed in explanted vitreous and in mouse eyes, including the immune inhibitor metalloproteases InhA1 and InhA2 [28,29]. A greater expression of InhA2 was detected in explanted vitreous compared to the levels detected in LB and BHI media. For InhA1, the expression in explanted vitreous was similar to InhA1 expression in BHI. A pangenome-wide study of ocular *Bacillus* isolates also reported molecular signatures of specific virulence factors, including the InhAs, which were strongly associated with this intraocular infection [30].

The InhA proteins are metalloproteases containing zinc-binding and catalytic active site residues similar to other metalloproteases such as PrtV of *Vibrio cholera*, thermolysin from *Bacillus thermopoteolyticus*, E-15 from *Serratia*, and elastase from *Pseudomonas aeruginosa* [31,32]. InhA1 is secreted by *Bacillus* during all phases of growth, and is associated with the exosporium [33]. The regulation of InhA1 is dependent on Spo0A, the key factor involved in the initiation of sporulation, and AbrB, which regulates sporulation gene expression [34,35]. InhA1 has been reported to hydrolyze the insect antibacterial proteins cecropin and attacin [36], degrade extracellular matrix proteins, and cleave fibronectin, laminin, and collagens types I and IV in tissue [37,38]. InhA1 also cleaves various exported proteins, including the protease NprA (Npr599 in *Bacillus anthracis)* [39]. Additionally, InhA1 contributes to *B. cereus* spore escape from macrophages [40,41]. Injection of purified *B. anthracis* InhA1 and nanoparticles conjugated to *B. anthracis* InhA1 into mice resulted in blood-brain barrier permeability, suggesting a potential role for InhA1 in meningitis [37].

More is known about InhA1 than about the other InhAs of *Bacillus*. InhA1 has a 66% protein identity to InhA2 and a 72% protein identity to InhA3. All three metalloproteases are secreted and contain a zinc-binding domain. InhA2 is involved in toxicity of *Galleria mellonella* after oral inoculation of spores, but InhA2 alone is not sufficient for virulence in this model [42,43]. In contrast to InhA1, InhA2 is regulated by PlcR and is repressed by Spo0A [25,43]. The transcription of *inhA3* is activated at the onset of sporulation by the quorum sensor NprR [44]. The specific functions of InhA2 and InhA3 in infection have not yet been described.

Due to the InhAs potential role in degrading important host tissue components and disrupting barriers, and the evidence that InhAs are expressed in an ocular infection-related environment, we hypothesized that the InhAs are involved in the *Bacillus* endophthalmitis pathogenesis. We used a well-characterized experimental model of endophthalmitis in mice to mimic human infection. Our study demonstrated that the absence of all three InhAs (InhA1, InhA2, and InhA3) together significantly reduced *Bacillus* virulence during ocular infection. Better knowledge of the underlying mechanisms of these virulence factors in the eye could lead to the identification of possible therapeutic targets that prevent vision loss in endophthalmitis patients.

## Results

### Absence of InhA1 in *Bacillus* Alters Growth and Proteolysis

The phenotypes of *B. thuringiensis* 407 (WT) and its isogenic InhA1-deficient mutant (Δ*inhA1*) were compared. WT and Δ*inhA1 in vitro* growth were compared by subculturing overnight cultures into fresh brain heart infusion (BHI) broth and quantifying every 2 hours. Figure 1A demonstrates that Δ*inhA1* had higher bacterial concentrations at 2, 4, and 6 hours compared to that of the WT strain, starting as early as 2 hours (P = 0.0263, 0.0065, 0.0059, respectively). Both strains reached similar concentrations at stationary phase at 8 hours. However, the overall growth rates were not different between the two strains (P = 0.2500, Figure 1B), suggesting that Δ*inhA1* entered exponential phase earlier than WT. In Figure 1C, hemolytic titers of 18 hour WT and Δ*inhA1* supernatants were similar (P ≥ 0.8678). Figure 1D shows the comparison of supernatant cytotoxicity of WT and Δ*inhA1* on human retinal pigment epithelial cells (RPEs). The strains had similar cytotoxicity (P = 0.0700). Proteolysis of WT and Δ*inhA1* on skim milk agar plates was also compared (Figure 1E). Clear lytic zone sizes around colonies were significantly different, with the Δ*inhA1 B. thuringiensis* exhibiting smaller proteolytic zones (P = 0.0006). Together, these results suggested that an absence of InhA1 affected bacterial growth and proteolytic activity, but not hemolysis or cytotoxicity.

**Figure 1.**
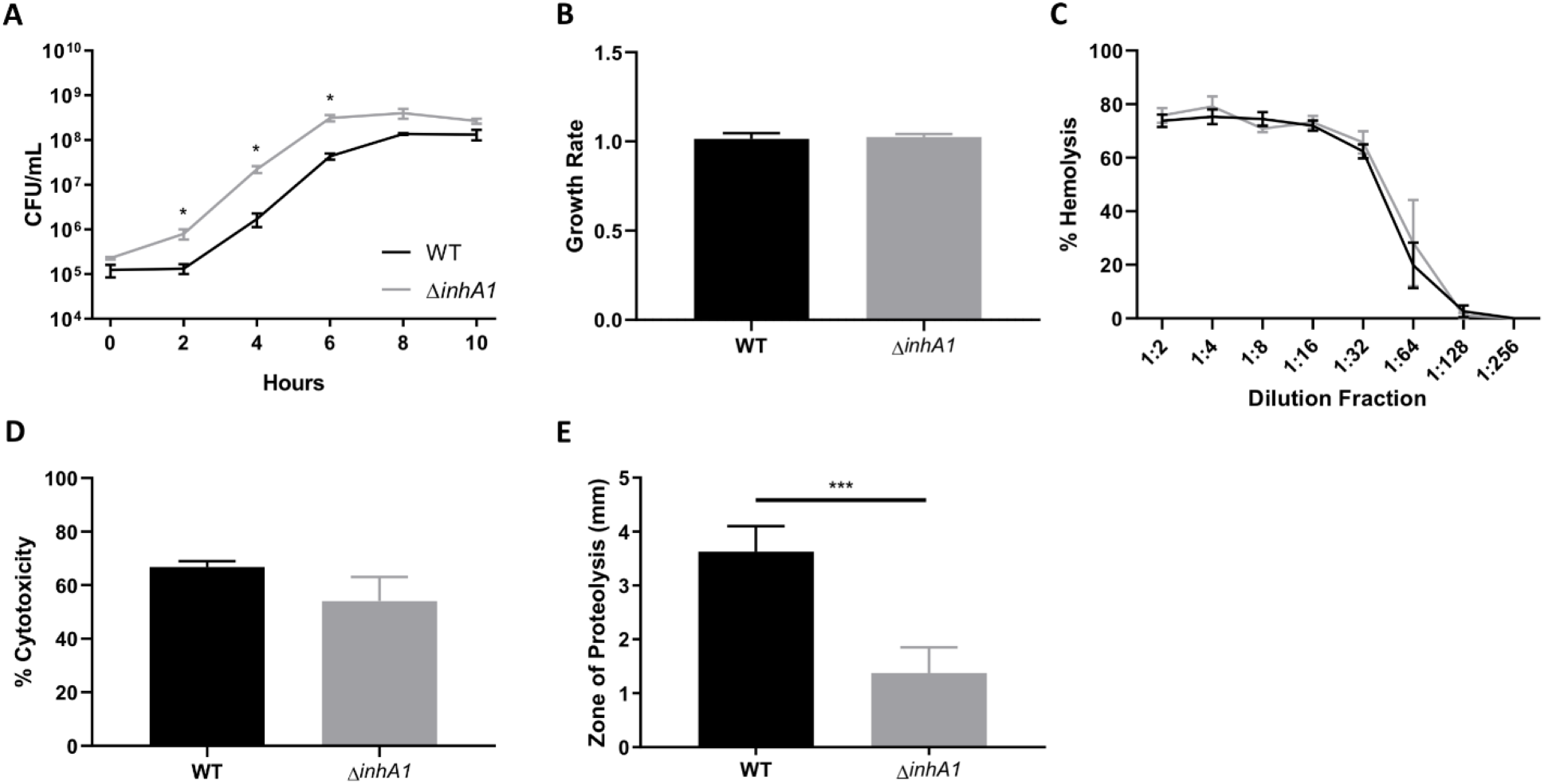
Absence of InhA1 in *Bacillus* alters growth and proteolysis. (**A**) *In vitro* growth curve of WT *B. thuringiensis* and its isogenic InhA1-deficient mutant (Δ*inhA1*) in BHI broth. CFU of Δ*inhA1 B. thuringiensis* were increased compared to WT at 2, 4, and 6 hours (*P* < 0.05). Values represent the mean ± SEM for *N* = 3 samples per time point. (**B**) Filter sterilized supernatants of WT and Δ*inhA1 B. thuringiensis* were compared for their hemolytic activities at varying dilutions (*P* > 0.05). (**C**) Cytotoxicity of filter sterilized overnight supernatants from WT and Δ*inhA1 B. thuringiensis* in human retinal pigment epithelial cells. No significant difference was observed in the cytotoxicity of these strains (*P* = 0.0744). Data represents the mean ± SEM of percent of cytotoxicity for *N* = 3 samples. (**D**) Proteolysis of WT and Δ*inhA1 B. thuringiensis* was compared by measuring lytic zones around colonies on milk agar. Lytic zones of Δ*inhA1 B. thuringiensis* were smaller compared to WT (P < 0.0005). Values represent the mean ± SEM for *N* = 4 samples.

### Absence of InhA1 Affects Intraocular Bacterial Burden But Not Inflammation in Endophthalmitis

The intraocular growth of WT and Δ*inhA1 B. thuringiensis* was quantified in the eyes of C57BL/6J mice (Figure 2). Mouse eyes were infected with approximately 200 CFU/eye of either WT or Δ*inhA1 B. thuringiensis*. At 6, 8, 10, and 12 hours postinfection, eyes were harvested and homogenized. Homogenates were plated on BHI agar and colonies were counted to quantify intraocular concentration (Figure 2A). Myeloperoxidase (MPO) concentrations were also quantified in these homogenates by ELISA (Figure 2B). Intraocular concentrations of WT and Δ*inhA1 B. thuringiensis* were significantly different at 6, 8, and 12 hours (P = 0.0164, 0.0359, 0.0332, respectively). The *in vitro* growth differences of WT and Δ*inhA1 B. thuringiensis* were reflected *in vivo*. MPO concentrations in the eyes infected with each strain were similar (P ≥ 0.0829), suggesting similar levels of inflammation. Overall, these results suggested that despite better growth of the Δ*inhA1* mutant, infection-related changes in eyes infected with either strain should be similar.

**Figure 2.**
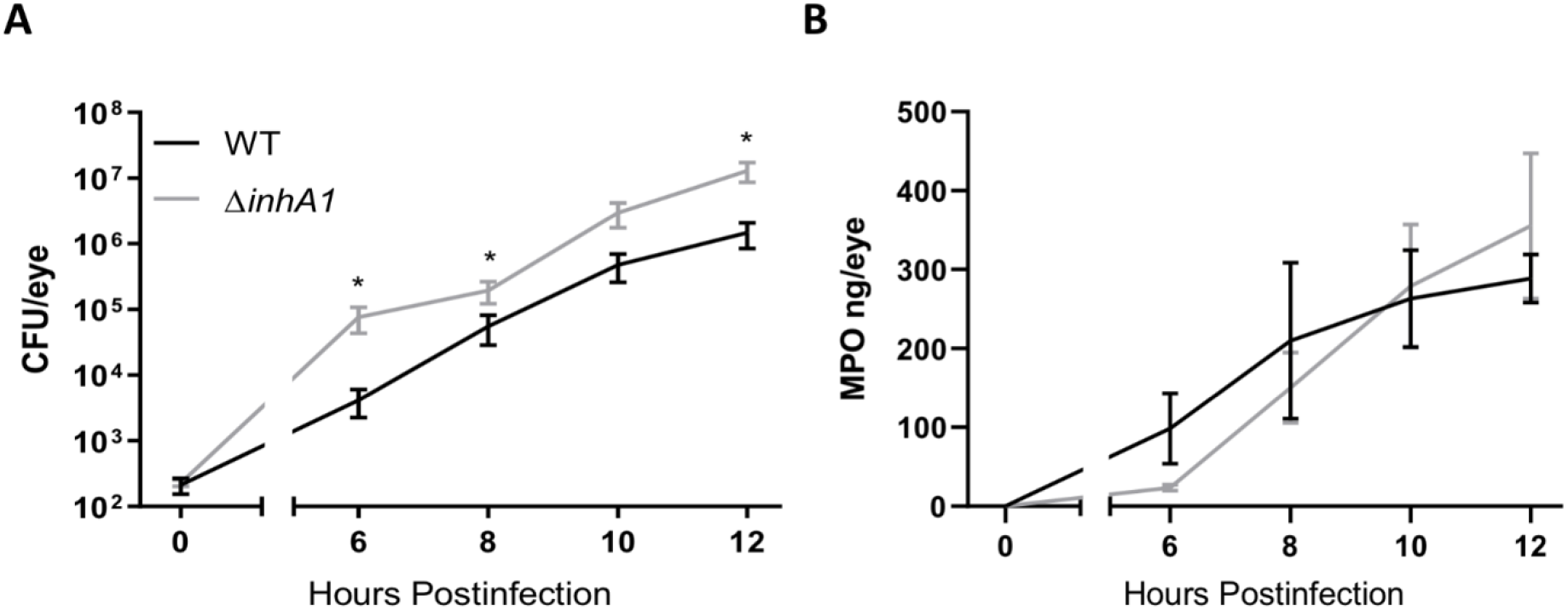
Absence of InhA1 Affects Intraocular Bacterial Burden But Not Inflammation in Endophthalmitis. C57BL/6J mouse eyes were injected with 200 CFU WT *B. thuringiensis* or its isogenic InhA1-deficient mutant (Δ*inhA1*). (**A**) At the indicated times postinfection, eyes were harvested and CFU quantified for bacterial intraocular growth. Data represents the mean ± SEM of log_10_ CFU/eye of *N* ≥ 4 eyes per time point for at least two separate experiments. ns: *P* > 0.05 at 0 and 10 hours postinfection. \**P* < 0.05 at 6, 8, and 12 hours postinfection. (**B**) Infected eyes were harvested and infiltration of PMN was assessed by quantifying MPO in whole eyes by sandwich ELISA. MPO levels of Δ*inhA1*-infected eyes were similar to WT strains at all time points. Values represent the mean ± SEM of MPO (ng/eye) of *N* ≥ 4 per time point for at least two separate experiments.

### Retinal Function is Not Preserved in the Absence of InhA1

To determine if the absence of InhA1 altered retinal damage in endophthalmitis, we analyzed WT- and Δ*inhA1*-infected mouse eyes using electroretinography (ERG). Figure 3 depicts retained A- and B-wave function and representative waveforms of infected eyes after 6, 8, 10, and 12 hours postinfection. The amplitude data indicated that retinal function in eyes infected with WT and Δ*inhA1 B. thuringiensis* was similar from 6 to 10 hours postinfection (P ≥ 0.2767, Figure 3A and Figure 3B). The function of retinal photoreceptor cells is represented by the A-wave function. Eyes infected with WT and Δ*inhA1 B. thuringiensis* showed similar reductions in A-wave function until 12 hours postinfection (P ≥ 0.2767, Figure 3A). The B-wave represents the function of rod bipolar cells, Muller cells, and second order neurons. At all time points, the B-wave function rapidly decreased in both WT and Δ*inhA1 B. thuringiensis*-infected eyes (P ≥ 0.3022, Figure 3B). The A-wave and B-wave retention responses declined to approximately 25% and 40% in eyes infected with WT or Δ*inhA1 B. thuringiensis*, respectively, after 12 hours. Figure 3C shows representative waveforms, which demonstrate the rapid decrease of retinal function in both A- and B-waves at 12 hours postinfection. These results demonstrated that the retinal function of eyes infected with WT or Δ*inhA1 B. thuringiensis* were similar, suggesting that the absence of InhA1 did not alter retinal function loss during experimental endophthalmitis.

**Figure 3.**
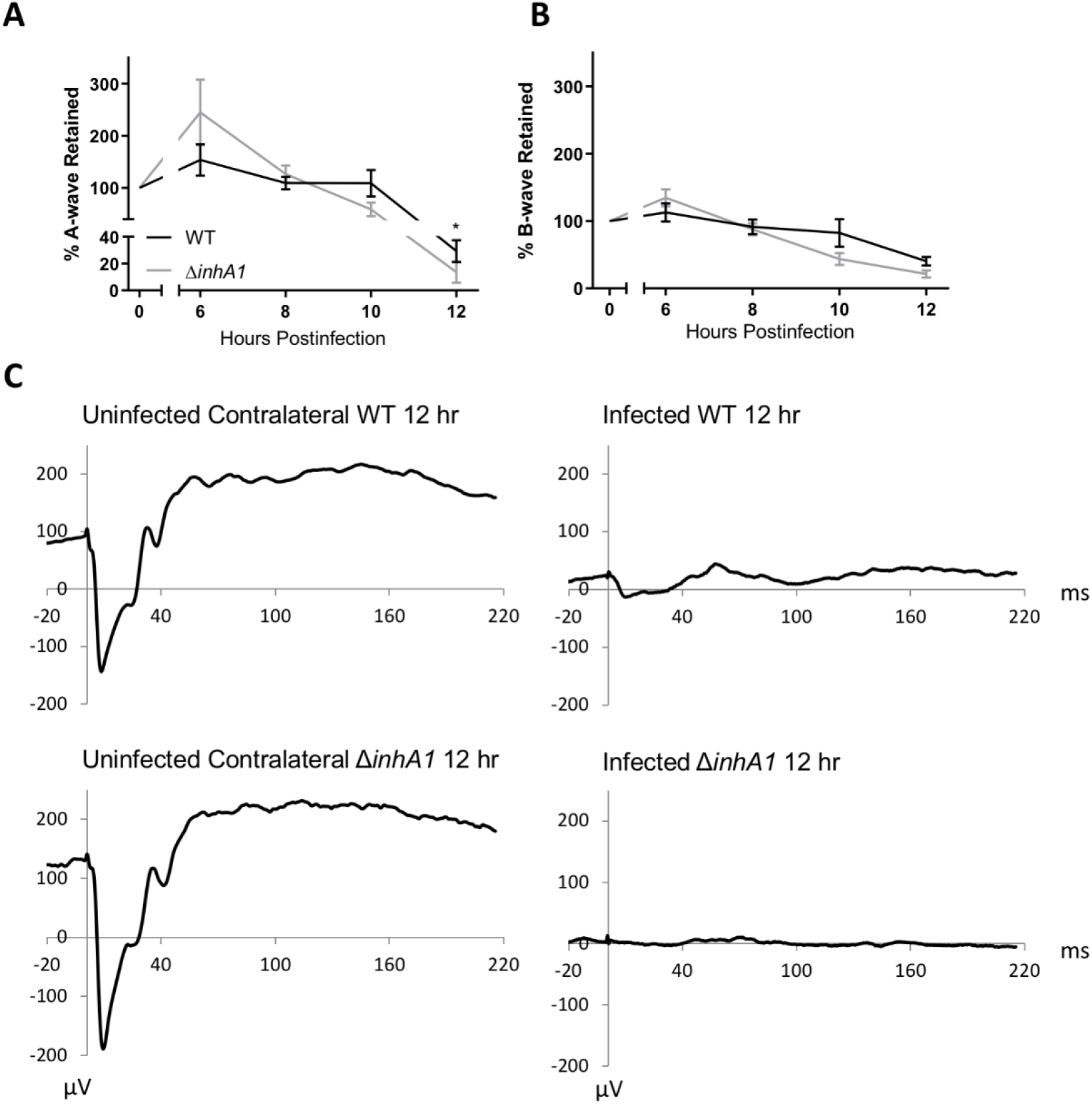
Retinal Function is Not Preserved in the Absence of InhA1. C57BL/6J mouse eyes were injected with 200 CFU WT or Δ*inhA1 B. thuringiensis* and retinal function was assessed by ERG. (**A**) Retained A-wave function of WT-infected eyes was similar to eyes infected with Δ*inhA2 B. thuringiensis* at 6, 8, and 10 hours postinfection (*P* > 0.05). \**P* < 0.05 at 12 hours postinfection. (**B**) B-wave function was also similar in eyes infected with WT and Δ*inhA1 B. thuringiensis* at 6, 8, 10, and 12 hours postinfection (*P* > 0.05). (**C**) Representative waveforms from eyes infected with WT or Δ*inhA1 B. thuringiensis* at 12 hours postinfection. In these mice, one eye was infected and the contralateral eye served as the uninfected control. Values represent the mean ± SEM of percentage amplitude retained per time point for at least two separate experiments. Data are representative of *N* ≥ 6 eyes per time point.

### Absence of InhA1 Does Not Preserve Ocular Architecture

The WT and Δ*inhA1 B. thuringiensis*-infected eyes were harvested and fixed, sectioned, and stained with hematoxylin and eosin (Figure 4). At all time points, the ocular architecture in both WT and Δ*inhA1 B. thuringiensis*-infected eyes were similar. Beginning at 8 hours postinfection, inflammatory cells entered the vitreous. At 10 hours postinfection, a significant amount of fibrin and inflammatory cells were observed in the vitreous. At 12 hours postinfection, severe inflammation, retinal detachment, and indistinguishable retinal layers were observed in the posterior segments of both WT and Δ*inhA1 B. thuringiensis*-infected eyes. These results showed that an absence of InhA1 did not reduce the damage observed in *Bacillus* endophthalmitis. This further suggests that InhA1 alone did not contribute to the pathogenesis of *Bacillus* endophthalmitis.

**Figure 4.**
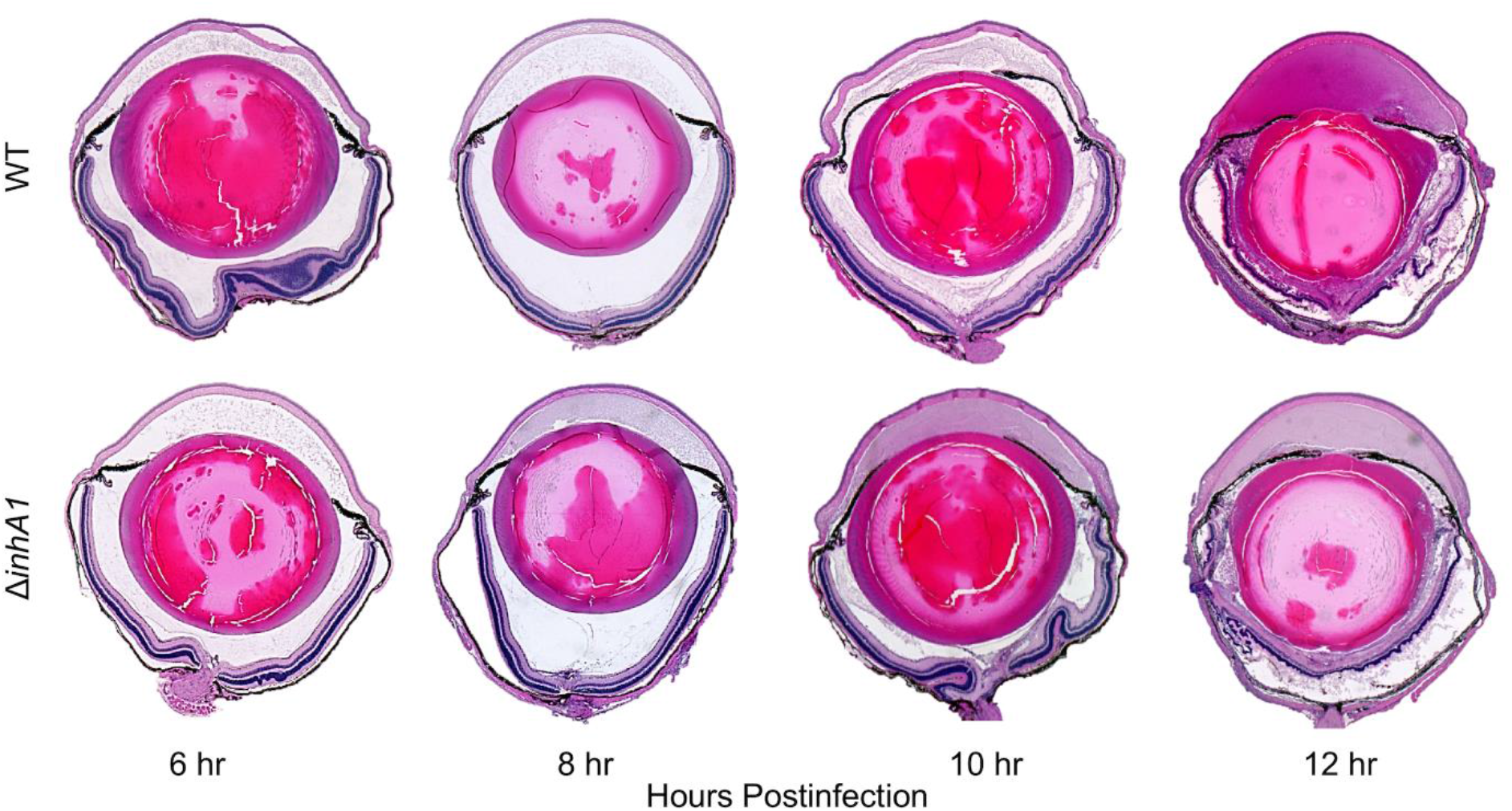
Absence of InhA1 Does Not Preserve Ocular Architecture. C57BL/6J mouse eyes were infected with 200 CFU of WT or Δ*inhA1 B. thuringiensis*. Infected eyes were harvested at 6, 8, 10, and 12 hours postinfection and processed for H&E staining. Magnification, ×10.

### Absence of InhA2 in *Bacillus* Alters Growth

Because InhA2 was also expressed in BHI, Luria-Bertani (LB) broth, and in *Bacillus* endophthalmitis-related environments [28–30], we also explored the contribution of this potential virulence factor. The growth and phenotypes of *B. thuringiensis* 407 (WT) and its isogenic InhA2-deficient mutant (Δ*inhA2*) were compared. Δ*inhA2* had higher bacterial concentrations than WT at 2, 6, and 8 hours (P = 0.0311, 0.0160, and 0.0075, respectively) (Figure 5A). Figure 5B demonstrates that although the InhA2 mutant and WT had similar growth rates, the InhA2 mutant reached stationary phase 2 hours before the WT B. *thuringiensis*. Δ*inhA2* also entered log phase 2 hours before the WT strain. Figure 5C showed no significant differences in the hemolytic activity of WT and Δ*inhA2 B. thuringiensis* supernatants from 18 hour cultures (P ≥ 0.9623). Cytotoxicity of human RPE was also similar between WT and Δ*inhA2 B. thuringiensis* supernatants (P = 0.7931, Figure 5D), as was proteolytic activity (P = 0.1359, Figure 5E). These results indicated that absence of InhA2 affected *in vitro* bacterial growth, but not proteolysis, hemolysis, or cytotoxicity. These results suggested that the intraocular growth of *Bacillus* lacking InhA2 might infect the mouse eye in a manner similar to that of the InhA1 mutant.

**Figure 5.**
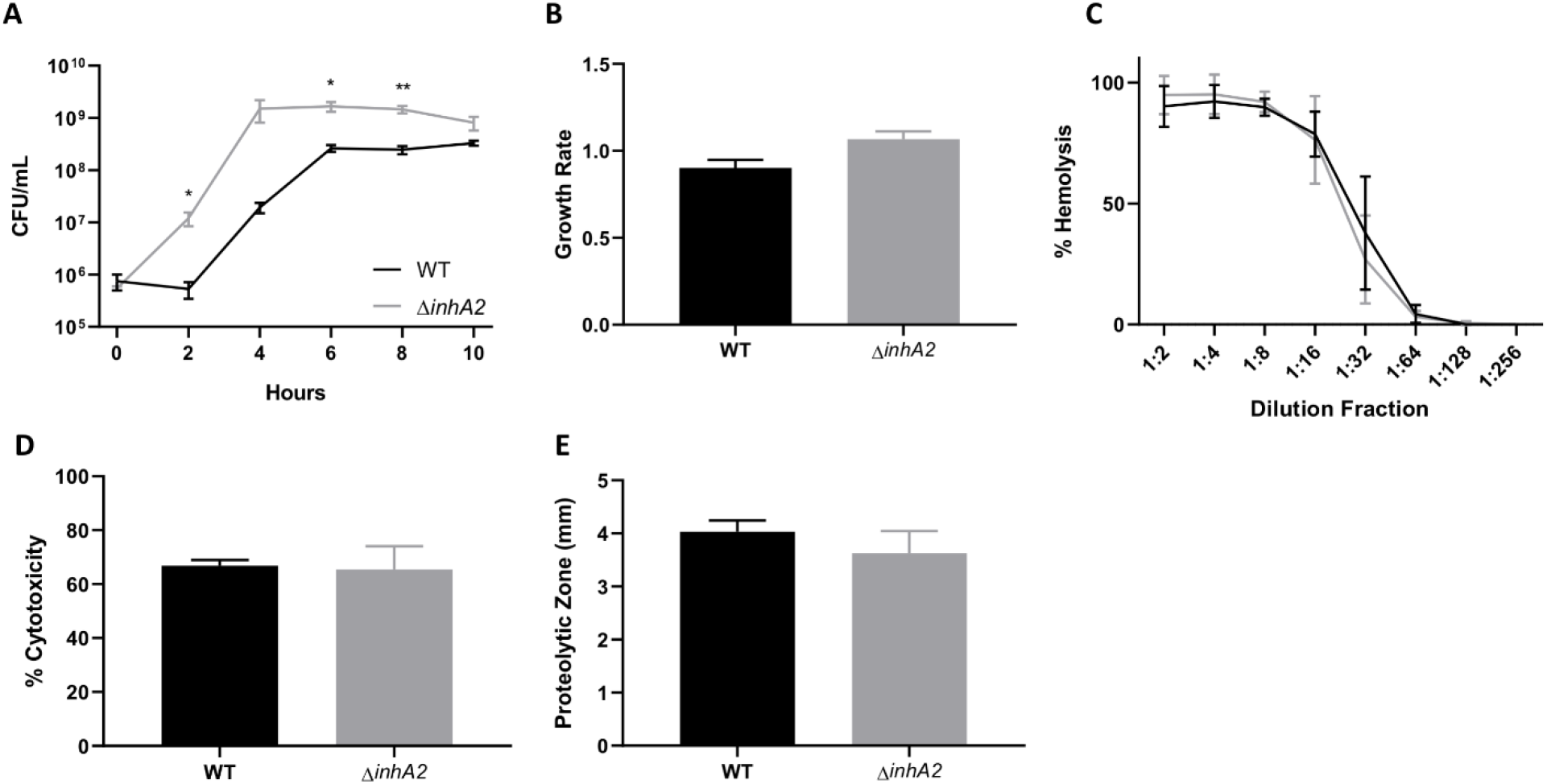
Absence of InhA2 in *Bacillus* Alters Growth. (**A**) *In vitro* growth curve of WT *B. thuringiensis* and its isogenic InhA2-deficient mutant (Δ*inhA2*) in BHI broth. CFU of Δ*inhA2 B. thuringiensis* were increased compared to WT at 2, 6, and 8 hours (*P* < 0.05). Values represent the mean ± SEM for *N* = 3 samples per time point. (**B**) Filter sterilized supernatants of WT and Δ*inhA2 B. thuringiensis* were compared for their hemolytic activities at varying dilutions (*P* > 0.05). (**C**) Cytotoxicity of filter sterilized overnight supernatants from WT and Δ*inhA2 B. thuringiensis* in human retinal pigment epithelial cells. No significant difference was observed in the cytotoxicity of these strains (*P* >0.05). Data represents the mean ± SEM of percent of cytotoxicity for *N* = 3 samples. (**D**) Proteolysis of WT and Δ*inhA2 B. thuringiensis* were compared by measuring lytic zones around colonies on milk agar. Lytic zones of Δ*inhA2 B. thuringiensis were similar to WT*(*P* > 0.05). Values represent the mean ± SEM for *N* = 4 samples.

### Absence of InhA2 Increases Bacterial Burden But Not Inflammation in Endophthalmitis

To determine if InhA2 contributed to intraocular growth and inflammation, the concentrations of bacteria and MPO were determined after 200 CFU of WT or Δ*inhA2 B. thuringiensis* were intravitreally injected into the eyes of mice. Figure 6A depicts the intraocular growth of WT and Δ*inhA2 B. thuringiensis* at 6, 8, 10, and 12 hours postinfection. The Δ*inhA2 B. thuringiensis*-infected eyes contained significantly more bacteria than WT-infected eyes at 6 and 12 hours postinfection (P = 0.0286 and 0.0087, respectively). Figure 6B shows that the intraocular MPO concentration was similar between eyes infected with WT or Δ*inhA2 B. thuringiensis* (P ≥ 0.4480). These results confirmed that an absence of InhA2 reflected the bacterial growth observed *in vitro* as well as the intraocular growth of the Δ*inhA1 B. thuringiensis*. These similarities suggested that the retinal changes in Δ*inhA2 B. thuringiensis*-infected eyes might be similar to the retinal changes observed in Δ*inhA1 B. thuringiensis*-infected eyes (Figure 2).

**Figure 6.**
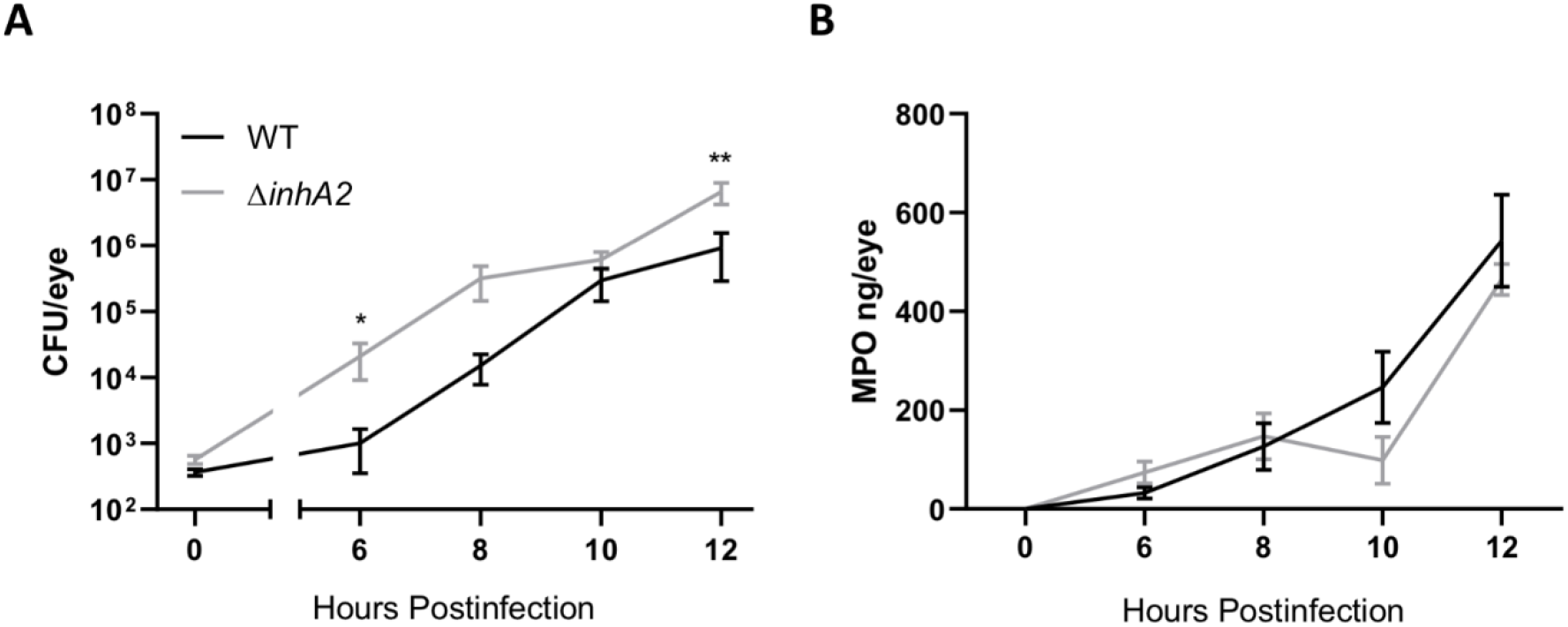
Absence of InhA2 Increases Bacterial Burden but not Inflammation in Endophthalmitis. C57BL/6J mouse eyes were injected with 200 CFU WT *B. thuringiensis* or its isogenic InhA2-deficient mutant (Δ*inhA2*). (**A**) At the indicated times postinfection, eyes were harvested and CFU quantified for bacterial intraocular growth. Data represents the mean ± SEM of log_10_ CFU/eye of *N* ≥ 4 eyes per time point for at least two separate experiments. ns: *P* > 0.05 at 0, 8, and 10 hours postinfection. \**P* < 0.05 at 6 hours postinfection, and **P < 0.005 at 12 hours postinfection. (**B**) Infected eyes were harvested and infiltration of PMN was assessed by quantifying MPO in whole eyes by sandwich ELISA. MPO levels of Δ*inhA2*-infected eyes were similar to WT strains at all time points. Values represent the mean ± SEM of MPO (ng/eye) of *N* ≥ 4 per time point for at least two separate experiments.

### Absence of InhA2 Does Not Affect Retinal Function

Retinal function was analyzed in WT and Δ*inhA2 B. thuringiensis*-infected mouse eyes (Figure 7). ERG data demonstrated that the A- and B-wave amplitudes of WT and Δ*inhA1 B. thuringiensis*-infected eyes were similar from 6 to 12 hours postinfection (P ≥ 0.3660, Figures 7A and 7B). Both the A- and B-wave retention responses rapidly decreased in WT and Δ*inhA2 B. thuringiensis*-infected eyes. Figure 7C shows the similarities in representative waveforms of eyes infected with these strains. This observation highlights the rapid decrease of retinal function in eyes infected with either WT or Δ*inhA2 B. thuringiensis*. Eyes infected with Δ*inhA2 B. thuringiensis* had retinal function loss similar to that of Δ*inhA1*-infected and WT-infected eyes, suggesting that the absence of InhA1 or InhA2 alone did not affect retinal function loss during *Bacillus* endophthalmitis.

**Figure 7.**
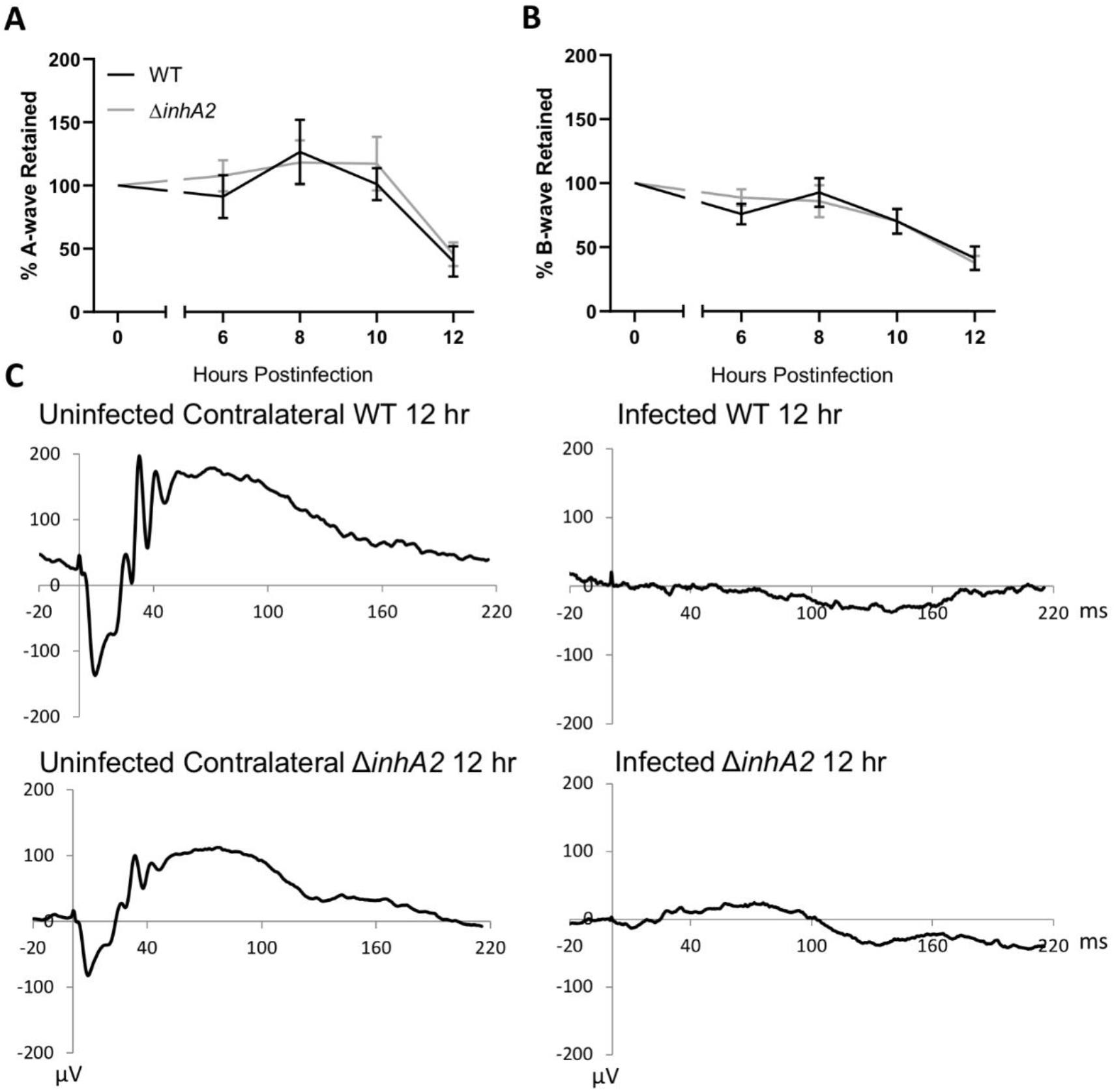
Absence of InhA2 Does Not Affect Retinal Function. C57BL/6J mouse eyes were injected with 200 CFU WT or Δ*inhA2 B. thuringiensis* and retinal function was assessed by ERG. (**A**) Retained A-wave function of WT-infected eyes was similar to eyes infected with Δ*inhA2 B. thuringiensis* at 6, 8, 10, and 12 hours postinfection (*P* > 0.05). (**B**) B-wave function was also similar in eyes infected with WT and Δ*inhA2 B. thuringiensis* at 6, 8, 10, and 12 hours postinfection (*P* > 0.05). (**C**) Representative waveforms from eyes infected with WT or Δ*inhA2 B. thuringiensis* at 12 hours postinfection. In these mice, one eye was infected and the contralateral eye served as the uninfected control. Values represent the mean ± SEM of percentage amplitude retained per time point for at least two separate experiments. Data are representative of *N* ≥ 6 eyes per time point.

### Ocular Damage and Inflammation are Similar Between WT and Δ*inhA2* Strains

To determine if InhA2 contributed to retinal damage, eyes infected with WT or Δ*inhA2 B. thuringiensis* were harvested for histological analysis at 6, 8, 10, and 12 hours postinfection (Figure 8). At every time point, the ocular architecture in both WT and Δ*inhA2 B. thuringiensis*-infected eyes was similar. At 8 hours postinfection, both WT and Δ*inhA2 B. thuringiensis*-infected eyes were inflamed. After 12 hours postinfection, there was retinal detachment and deterioration of retinal layers in the posterior segment of both WT and Δ*inhA2 B. thuringiensis*-infected eyes. These results showed that the absence of InhA2 alone, like the absence of InhA1 alone, did not alter the ocular damage observed in *Bacillus* endophthalmitis.

**Figure 8.**
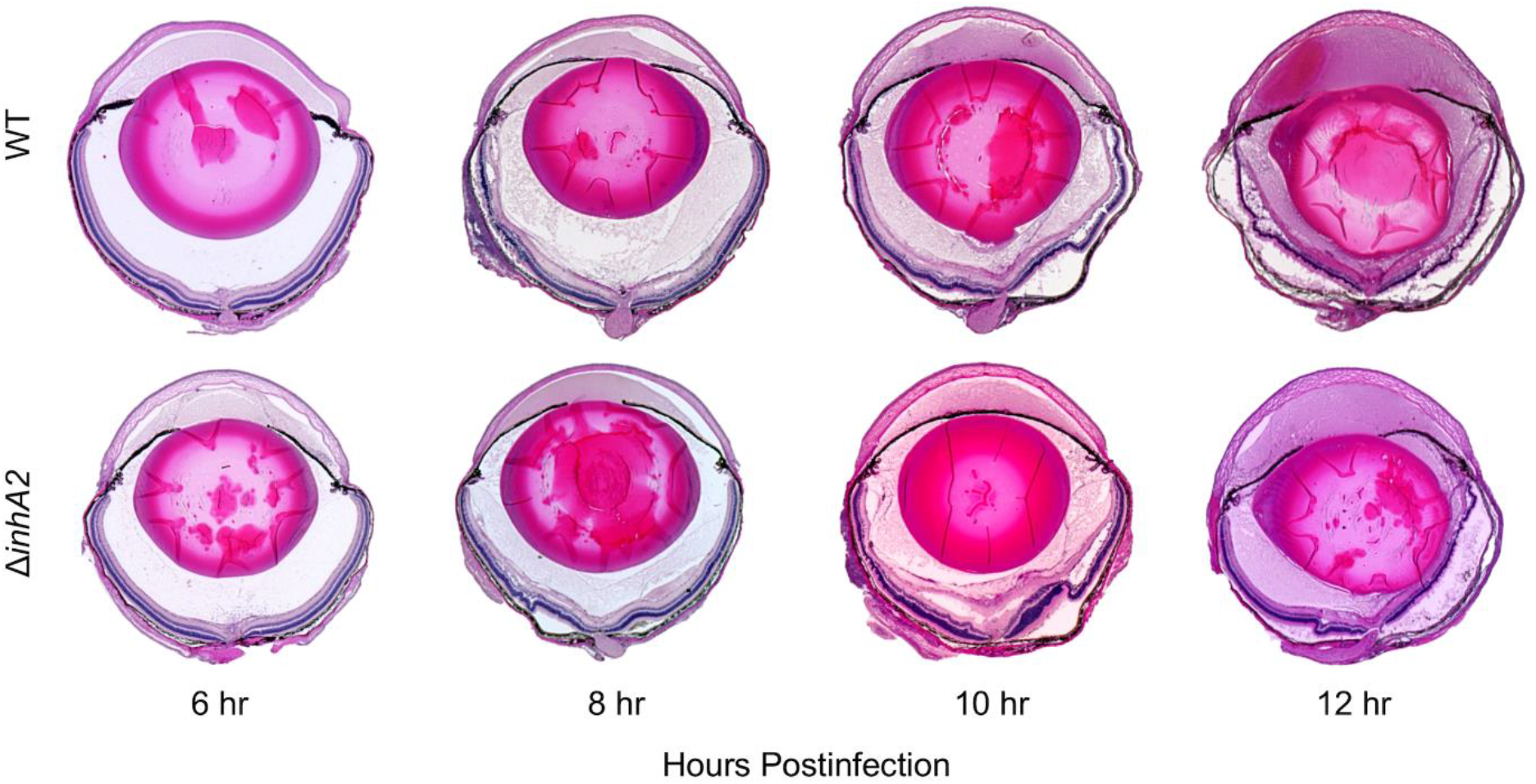
Ocular Damage and Inflammation are Similar Between WT and Δ*inhA2* Strains in Endophthalmitis. C57BL/6J mouse eyes were infected with 200 CFU of WT or Δ*inhA2 B. thuringiensis*. Infected eyes were harvested at 6, 8, 10, and 12 hours postinfection and processed for H&E staining. Magnification, ×10.

### Compensation of InhA Expression in the Single InhA Mutants

To understand the *in vitro* and *in vivo* results observed with the single InhA mutants, we sought to determine if the absence of one InhA affected the expression of other InhAs. Therefore, we analyzed the expression of *inhA1, inhA2, and inhA3* in Δ*inhA1* and Δ*inhA2 B. thuringiensis*. Additionally, the expression of all three *inhAs* were examined in Δ*inhA1-3 B. thuringiensis* as a negative control. Expression in all mutants was compared to that of WT (Figure 9). The expression of *inhA2* and *inhA3* in the Δ*inhA1* strain was elevated, but not statistically significantly different from that of WT (Ct values, P ≥ 0.3342). The expression of *inhA1* in the Δ*inhA2* strain was significantly greater than WT (Ct values, P = 0.0036). The expression of *inhA3* in the Δ*inhA2* strain was elevated, but not statistically significantly different from that of WT (Ct values, P = 0.1881). Overall, these results suggested that an absence of expression of a single *inhA* caused an elevated, although not always statistically significant, expression of the other *inhAs*. This potential compensation could explain why infections with Δ*inhA1* or Δ*inhA2* mutants were not different from WT infections, despite the differences in growth. To examine the role of InhAs as a whole in this disease, a *Bacillus* mutant lacking all three InhAs was tested in subsequent experiments.

**Figure 9.**
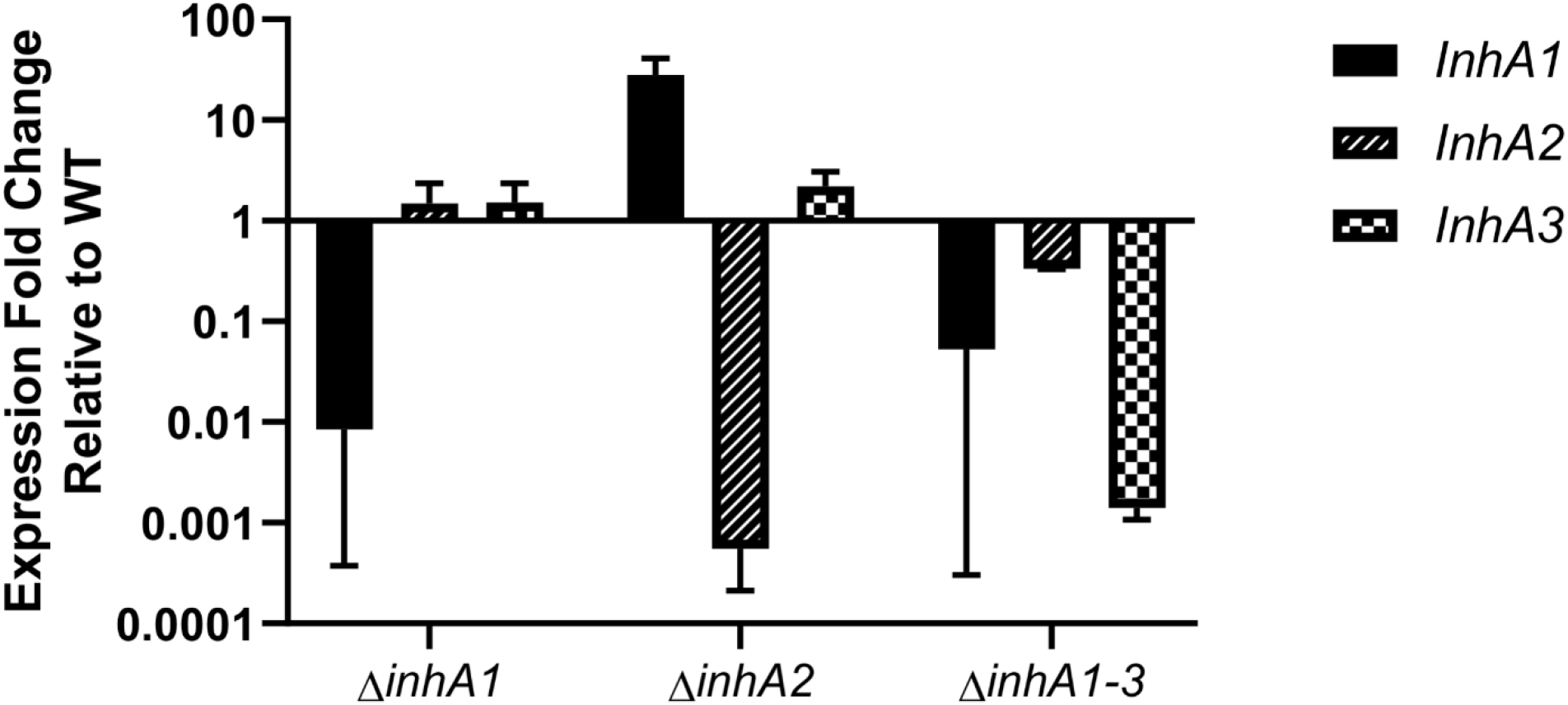
Compensation of InhA Expression in Single InhA Mutants. Quantitative RT-PCR of mutant strains detecting *inhA1, inhA2, and inhA3* in overnight cultures grown in BHI. 16S ribosomal RNA was used as a control. Values represent the mean ± SD of expression fold change relative to the expression in WT. Data are representative of at least two separate experiments, and are representative of N=3.

### Absence of InhA1, InhA2, and InhA3 in *Bacillus* Alters Proteolysis

To determine if all three *Bacillus* InhAs together were involved in the severity of *Bacillus* endophthalmitis, a strain lacking all three InhAs (Δ*InhA1-3*) was tested. Phenotypes of the Δ*inhA1-3* strain were compared to that of its WT *Bacillus* parent strain (Figure 10). Unlike the single InhA mutants, the *in vitro* growth of WT and Δ*inhA1-3* in BHI was similar at every time point (P ≥ 0.3586, Figure 10A). Figure 10B demonstrates that both strains had similar growth rates (P = 0.7220). Figure 10C shows similarities in hemolytic activities of the supernatants of Δ*inhA1-3* and the WT strain, except at the dilution of 1:32 (P = 0.0001). Both strains had similar cytotoxicity against RPE cells (P = 0.8250, Figure 10D). The proteolytic zones of Δ*inhA1-3 B. thuringiensis* colonies were significantly smaller compared to that of WT colonies (P = 0.0018, Figure 10E). These results showed that an absence of all the InhAs did not affect bacterial growth, hemolysis, or cytotoxicity, but reduced proteolysis. Overall, the results suggested that the Δ*InhA1-3* growth might be similar to WT *in vivo*.

**Figure 10.**
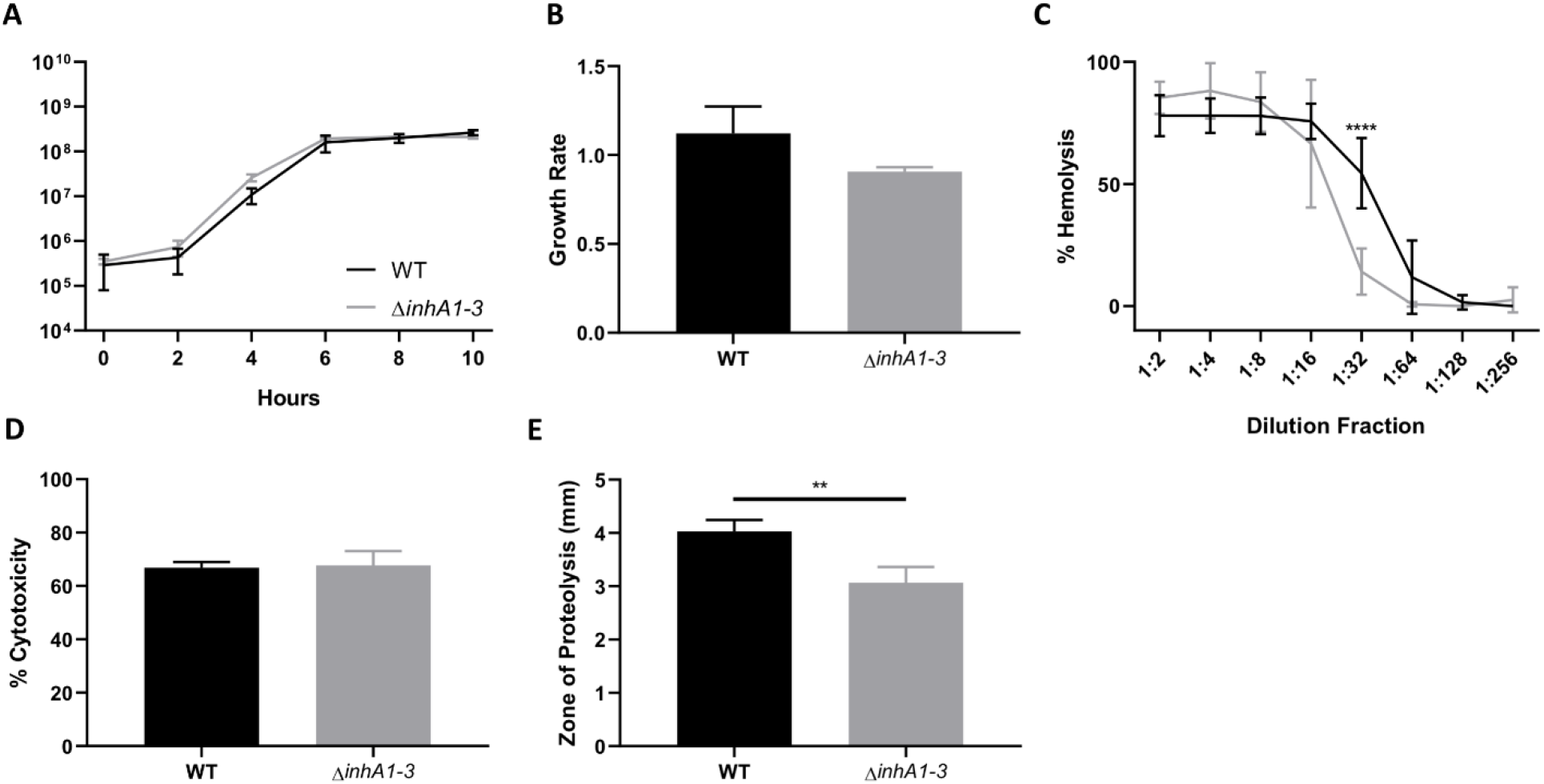
Absence of InhA1, InhA2, and InhA3 in *Bacillus* Alters Proteolysis. (**A**) *In vitro* growth curve of WT *B. thuringiensis* and its isogenic InhA1-3-deficient mutant (Δ*inhA1-3*) in BHI broth. CFU of Δ*inhA1-3 B. thuringiensis* was similar to WT at all time points (*P* < 0.05). Values represent the mean ± SEM for *N* = 3 samples per time point. (**B**) Filter sterilized supernatants of WT and Δ*inhA1-3 B. thuringiensis* were compared for their hemolytic activities at varying dilutions (*P* > 0.05). (**C**) Cytotoxicity of filter sterilized overnight supernatants from WT and Δ*inhA1-3 B. thuringiensis* in human retinal pigment epithelial cells. A significant difference was observed in the cytotoxicity of these strains at the 1:32 dilution fraction (*P* >0.05). Data represents the mean ± SEM of percent of cytotoxicity for *N* = 3 samples. (**D**) Proteolysis of WT and Δ*inhA1-3 B. thuringiensis* were compared by measuring lytic zones around colonies on milk agar. Lytic zones of Δ*inhA1-3 B. thuringiensis* were significantly less compared to WT (*P* > 0.05). Values represent the mean ± SEM for *N* = 4 samples.

### Absence of InhAs 1, 2, and 3 Alters Intraocular Bacterial Burden and Inflammation

Because preliminary data suggested that infections with the Δ*inhA1-3* would be less severe, intraocular growth and MPO concentrations were quantified in eyes infected with Δ*inhA1-3* after 12 hours. The intraocular bacterial concentrations of Δ*inhA1-3 B. thuringiensis* were lower than that of WT, with significant differences at 12, 14, and 16 hours postinfection (P = 0.0002, 0.0022, 0.0095, respectively, Figure 11A). At 16 hours postinfection, the concentrations Δ*inhA1-3 B. thuringiensis* were below the limit of detection in the eye. The MPO concentrations of Δ*inhA1-3*-infected eyes were also significantly lower compared to WT-infected eyes at 6, 10, and 14 hours postinfection (P = 0.0079, 0.0286, 0.0022, respectively, Figure 11B). These differences showed that an absence of all three InhAs resulted in a lower intraocular bacterial concentration that cleared approximately 16 hours postinfection. These differences suggested that the retinal function in Δ*inhA1-3 B. thuringiensis*-infected eyes should be preserved, compared with eyes infected with the WT strain.

**Figure 11.**
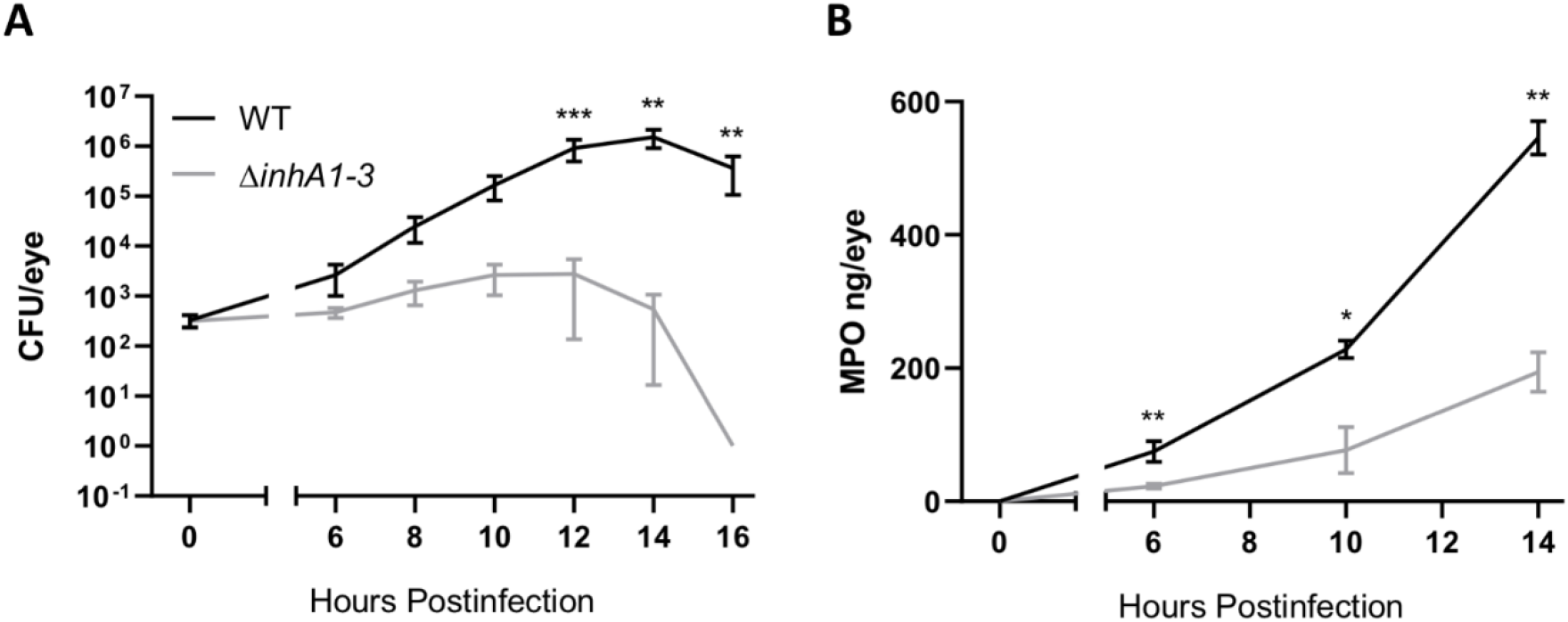
Absence of InhA1-3 Alters Intraocular Bacterial Burden and Inflammation in Endophthalmitis. C57BL/6J mouse eyes were injected with 200 CFU WT *B. thuringiensis* or its isogenic InhA1-3-deficient mutant (Δ*inhA1-3*). (**A**) At the indicated times postinfection, eyes were harvested and CFU quantified for bacterial intraocular growth. Data represents the mean ± SEM of log_10_ CFU/eye of *N* ≥ 4 eyes per time point for at least two separate experiments. P > 0.05 at 12, 14, and 16 hours postinfection. **P < 0.005 at 14 and 16 hours postinfection, and ***P < 0.0005 at 12 hours postinfection. (**B**) Infected eyes were harvested and infiltration of PMN was assessed by quantifying MPO in whole eyes by sandwich ELISA. MPO levels of Δ*inhA1-3*-infected eyes were significantly less compared to WT strains at all time points. *P < 0.05 at 10 hours postinfection, and **P < 0.005 at 6 and 14 hours postinfection. Values represent the mean ± SEM of MPO (ng/eye) of N ≥ 4 per time point for at least two separate experiments.

### Retained Retinal Function in Eyes Infected with *Bacillus* Lacking all InhAs

Retinal function analysis of mouse eyes infected with either WT or Δ*inhA1-3 B. thuringiensis* was performed to determine if the absence of the three InhAs affected retinal function. In Figure 12, ERGs depict retained retinal function in eyes infected with Δ*inhA1-3 B. thuringiensis*, which was significantly greater than that of WT-infected eyes at 12, 14, and 16 hours postinfection for A-wave (P ≤ 0.0358) and B-wave (P ≤ 0.0015). The photoreceptor function in Δ*inhA1-3 B. thuringiensis*-infected eyes was approximately 80% whereas WT-infected eyes had a photoreceptor function that decreased to approximately 40% at 16 hours postinfection (Figure 12A). Retinal function in Δ*inhA1-3 B. thuringiensis*-infected eyes was retained to approximately 60%, while function in the WT-infected eyes decreased to approximately 30% at 16 hours postinfection (Figure 12B). Figure 12C shows representative waveforms of WT and Δ*inhA1-3 B. thuringiensis*-infected eyes at 14 hours postinfection, which had significantly reduced amplitudes in WT-infected eyes and retained amplitudes in Δ*inhA1-3 B. thuringiensis*-infected eyes. This observation indicated that in the absence of InhA1, InhA2, and InhA3, retinal function during *Bacillus* endophthalmitis was preserved.

**Figure 12.**
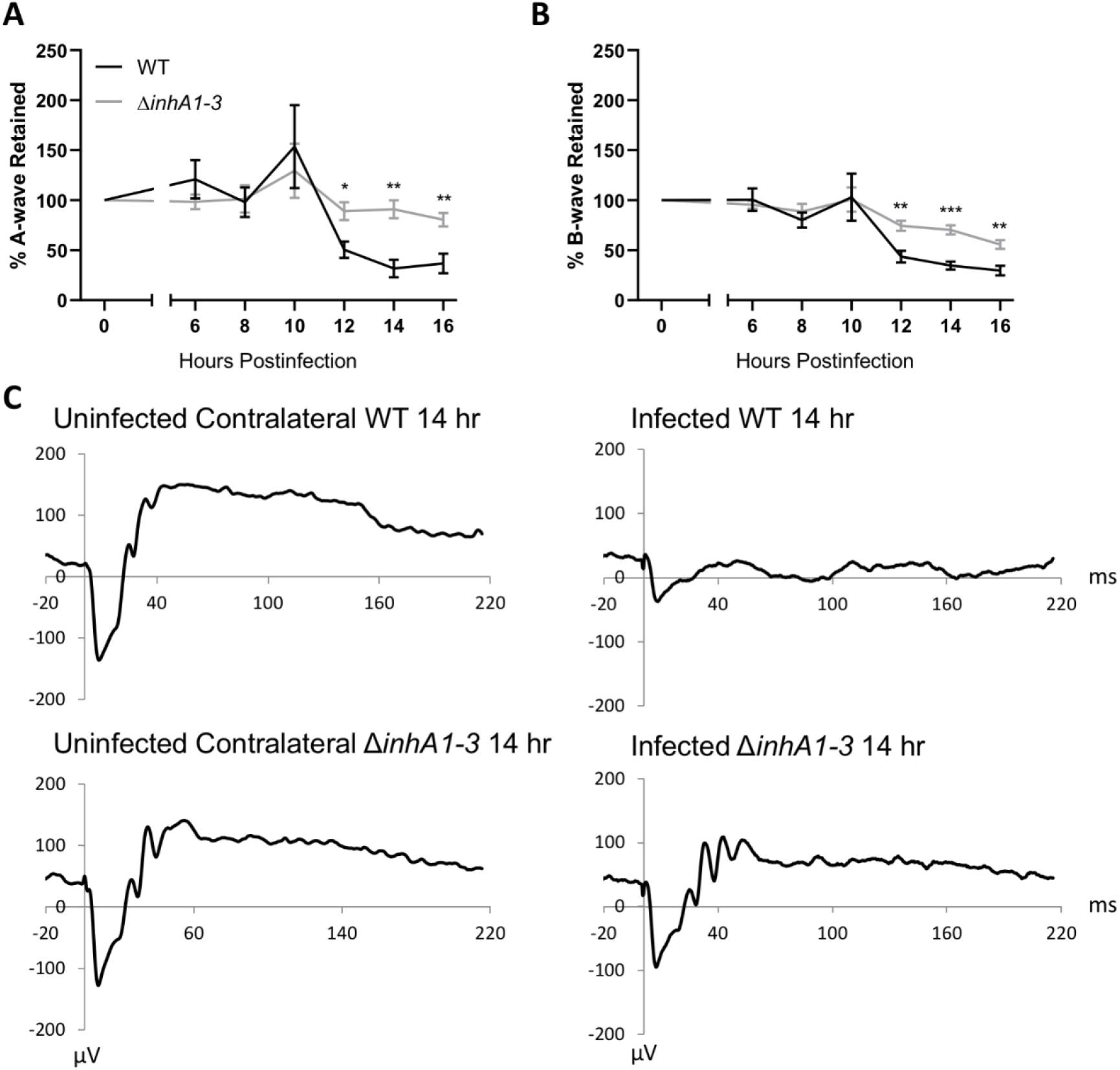
Retained Retinal Function in eyes infected with *Bacillus* lacking InhA1-3. C57BL/6J mouse eyes were injected with 200 CFU WT or Δ*inhA1-3 B. thuringiensis* and retinal function was assessed by ERG. (**A**) Retained A-wave function of WT-infected eyes was significantly higher to eyes infected with Δ*inhA1-3 B. thuringiensis* at 12, 14, and 16 hours postinfection. (**B**) B-wave function was also higher in eyes infected with Δ*inhA2 B. thuringiensis* at 12, 14, and 16 hours postinfection. *P < 0.05, **P < 0.005, and ***P < 0.0005. (**C**) Representative waveforms from eyes infected with WT or Δ*inhA1-3 B. thuringiensis* at 12 hours postinfection. In these mice, one eye was infected and the contralateral eye served as the uninfected control. Values represent the mean ± SEM of percentage amplitude retained per time point for at least two separate experiments. Data are representative of N ≥ 6 eyes per time point.

### Ocular Architecture is Preserved in the Absence of all InhAs

Histology of WT or Δ*inhA1-3 B. thuringiensis*-infected eyes is illustrated in Figure 13. Eyes were harvested for histology at 6, 8, 10, and 12 hours postinfection. Mouse eyes exhibited similar degrees of inflammation until 12 hours postinfection. At 12 hours postinfection, the WT eyes had retinal detachments and loss of retinal layers. However, the retinas of Δ*inhA1-3 B. thuringiensis*-infected eyes were attached, and the layers of the retina were distinguishable. The Δ*inhA1-3 B. thuringiensis*-infected eyes had fibrin and inflammatory cells present in the vitreous at 12 hours postinfection, but to a lesser degree than that of WT-infected eyes. Together, these results demonstrated that the absence of all three InhAs (InhA1, InhA2, and InhA3) significantly reduced *Bacillus* virulence during intraocular infection.

**Figure 13.**
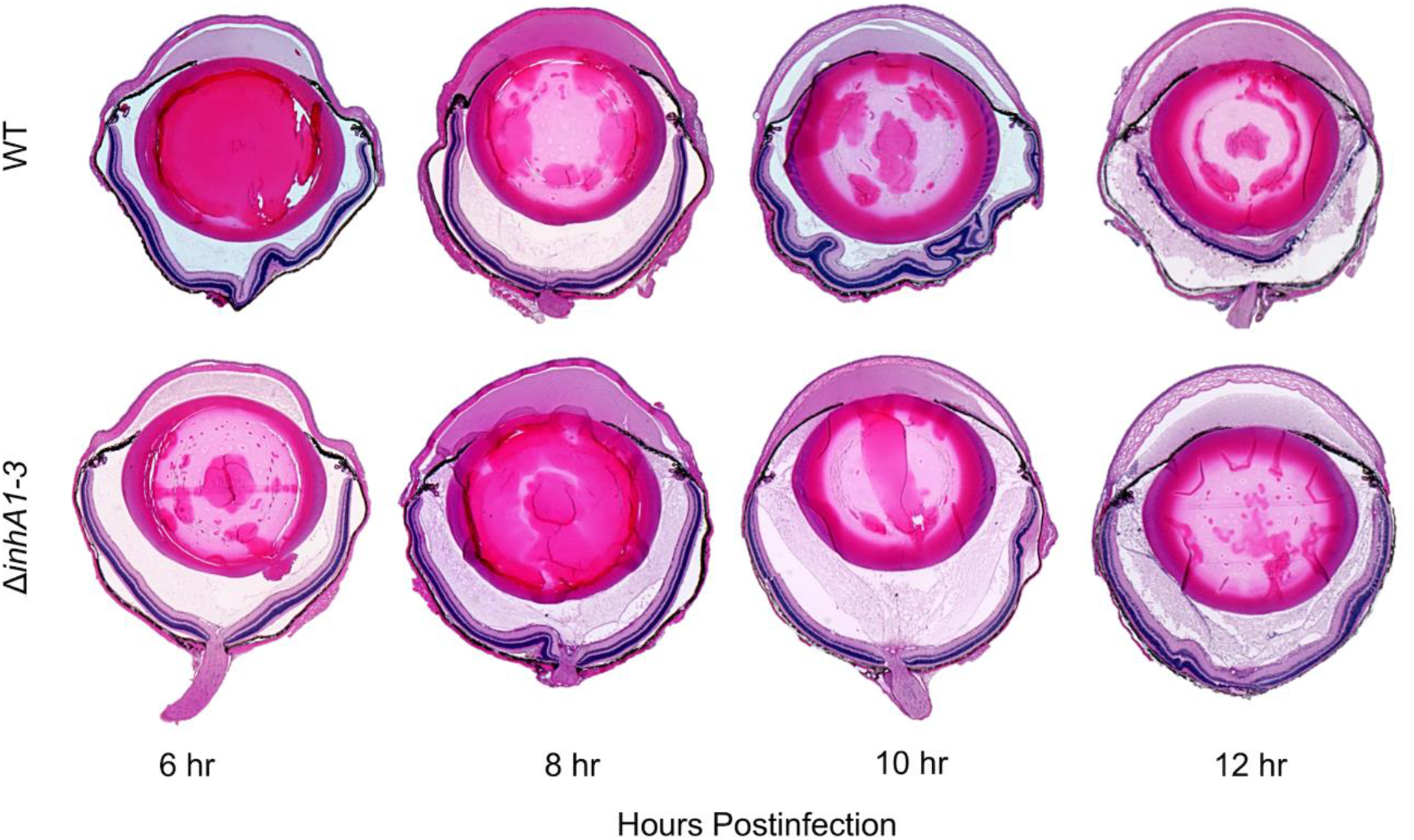
Ocular architecture is preserved in the absence of InhA1-3. C57BL/6J mouse eyes were infected with 200 CFU of WT or Δ*inhA1-3 B. thuringiensis*. Infected eyes were harvested at 6, 8, 10, and 12 hours postinfection and processed for H&E staining. Magnification, ×10.

## Discussion

*Bacillus* is capable of producing a rapid and severe intraocular infection, which often results in loss of vision or the eye itself. During experimental endophthalmitis, *Bacillus* induces an explosive inflammatory response with vascular permeability and PMN infiltration into the vitreous beginning at approximately 4 hours postinfection [1,6]. This blood-retinal barrier permeability and neutrophil infiltration into a normally immune privileged environment is detrimental to vision. The immune response in the eye is tightly regulated -- the vitreous is avascular, there is a lack of lymphatic vessels and antigen presenting cells, and immunosuppressive factors are present [45–47]. Bacterial endophthalmitis and other ocular diseases with inflammation compromise immune privilege. A combination of toxin activities, the early innate immune response, and subsequent ocular damage during endophthalmitis allows the components of blood from the retinal vasculature enter the vitreous [20,21,48–52]. The retinal vascular permeability and PMN infiltration observed in *Bacillus* endophthalmitis is due, in part, to *Bacillus* virulence factors and their effect on ocular barrier cells, such as the RPE [20,21]. Identifying virulence factors that contribute to processes that result in compromised ocular clarity are of interest in understanding how *Bacillus* induces a rapid and severe disease.

*Bacillus cereus sensu lato* group species produce many virulence factors that may contribute to ocular inflammation and damage during *Bacillus* endophthalmitis [12,17–20,23,26]. For *B. cereus* and *B. thuringiensis*, a majority of these virulence factors are controlled by the pleotropic regulator of extracellular virulence, PlcR. We reported that in the absence of PlcR-regulated toxin expression, *Bacillus* is capable of inducing inflammatory cell influx, blood retinal permeability, and retinal function loss in experimental endophthalmitis, albeit slower than that caused by WT *Bacillus* [19,21]. This suggests that virulence factors not regulated by PlcR contribute to intraocular infection as well.

To identify the specific virulence factors that contribute to *Bacillus* endophthalmitis, recent studies have observed molecular signatures and virulence factor gene expression in *Bacillus*-infected mouse eyes. In particular, the InhA metalloproteases of *Bacillus* are highly expressed in explanted vitreous [28] and are strongly associated with experimental *Bacillus* endophthalmitis [29,30]. The InhAs have a wide range of functions in many members of the *Bacillus cereus sensu lato* group. Originally identified for hydrolyzing insect antibacterial proteins [36], the InhA1 protein has now been demonstrated to be involved in autocleavage events which may contribute to *Bacillus*’ ability to escape from host macrophages and modulate its own secretome [40,53]. InhA1 has been observed to contribute to blood-brain barrier permeability by degrading ZO-1 [37], a tight junction protein found in retinal barriers [20,21]. These studies highlight the potential importance of InhAs in infection, so we sought to determine their contribution to the severity of *Bacillus* endophthalmitis.

To ensure that the absence of InhAs did not affect hemolysis and cytotoxicity, we compared these activities in supernatants of InhA mutant and WT *B. thuringiensis*. The absence of InhA1 affected proteolysis, but not hemolysis or cytotoxicity. This was not the case for InhA2, since Δ*inhA2 B. thuringiensis* was as proteolytic as WT. This result may have been attributed to the elevated expression of InhA1 and InhA3 in the Δ*inhA2* mutant compared to WT. Although InhA2 and InhA3 were expressed in the Δ*inhA1* mutant, the proteolysis of Δ*inhA1 B. thuringiensis* was significantly less than WT. InhA1 has been shown to be important for degrading tight junction proteins, plasma, and matrix proteins [37,38,54]. Thus, InhA1 may be important for retinal vascular permeability and intraocular tissue destruction via degradation of tight junction and extracellular matrix protein during *Bacillus* endophthalmitis.

In this study, we investigated the effects of *Bacillus thuringiensis* InhA mutations on intraocular growth, inflammation, and retinal function during experimental *Bacillus* endophthalmitis. Our findings showed that the absence of single InhAs affected the expression of the other InhAs, and that bacterial growth *in vitro* was affected by the absence of InhA1 or InhA2, resulting in higher concentrations of bacteria beginning at the exponential phase of growth. A similar growth phenotype was observed *in vivo* during *Bacillus* endophthalmitis. This growth phenotype may be a consequence of the expression of other InhAs when one InhA is absent. Another possible explanation for this difference is that InhAs might affect bacterial physiology. We inoculated fresh media with stationary phase *Bacillus*, so *Bacillus* must re-enter the logarithmic state before multiplying. It is possible that the physiological state of the Δ*inhA1* and Δ*inhA2* strains facilitated a faster switch to logarithmic phase after inoculation in fresh medium. The InhA metalloproteases are a part of the M6 evolutionary protein family, which has been identified in environmental genera such as *Clostridioides, Geobacillus, Shewanella*, and *Vibrio* [55]. This type of protease might contribute to environmental persistence, nutrient acquisition, and survival. In a study investigating bacterial zinc metalloproteases from *Aeromonas salmonicida*, protease-deficient *A. almonicida* strains had slower rates of bacterial growth in media with heat-inactivated serum [56]. In untreated serum, the growth rates were quickly reduced in these mutants, suggesting that proteases play a critical role in the early stages of infection process by protecting bacteria against complement-mediated killing or other serum bactericidal effects. This also suggested that proteolytic activity might provide nutrients for continued growth and proliferation. A similar phenomenon may be at play in this study, in which a mutant lacking all three InhAs is unable to provide nutrients in the vitreous for patterns of growth similar to wild type, leading to clearance starting at 12 hours postinfection. The Δ*inhA1-3* strain may have grown similar to WT *in vitro* because of the readily available nutrients in the BHI medium. The growth effects in the single mutants may be explained by the availability of nutrients via expression of the other InhAs. *Bacillus* may utilize these InhAs to prolong survival in the ocular environment by degrading proteins, such as collagen fibers of types II, V, IX and XI in the vitreous. In the absence of the three InhAs, *Bacillus* may not have been able to degrade these proteins, and the lack of nutrient availability resulted in a lower burden that was more easily cleared.

The inflammatory response in eyes infected with Δ*inhA1* or Δ*inhA2* was similar to that seen in WT-infected eyes. These observations were unexpected considering the increased intraocular growth of the single mutants compared to WT. In eyes infected with *Bacillus* lacking InhAs 1, 2, and 3, the inflammatory response was delayed. These observations may have been due to the compensating expression of the other InhAs in the single mutants, which may have facilitated growth to concentrations which triggered the activation of innate immune pathways, blood-retinal barrier permeability, and PMN infiltration [57,58]. This effect may have been similar to how *B. anthracis* InhA contributes to breakdown of blood-brain barriers via degradation of ZO-1, leading to inflammatory cell infiltration and hemorrhaging in the mouse brain [37]. We reported that *B. cereus*-infected eyes have little to no expression of ZO-1 in the RPE at 12 hours postinfection [21]. Also, when injected into mouse eyes, WT *B. cereus* and Δ*plcR B. cereus* supernatants induced permeability of the blood ocular barrier [21]. Therefore, secreted InhAs may contribute to the degrading of ZO-1 and permeability of the blood-retinal barriers, leading to infiltration of PMNs into the vitreous. PMN are the first and most abundant inflammatory cells infiltrating into the eye during *B. cereus* endophthalmitis [1,6]. These cells are capable of phagocytosing *B. cereus in vitro* [57]. InhA1 has been shown to be important for allowing *B. cereus* to escape from macrophages [40]. Whether the InhAs are important in protecting *Bacillus* from PMN-mediated killing in the eye is an open question.

Both bacteria and the inflammatory response contribute to retinal damage. Damage to retinal cells and tissues may result in substantial and permanent vision loss, which is typical for *Bacillus* endophthalmitis [9–11]. Because the expression of InhA has been associated with *Bacillus* ocular infection in mice, and in toxicity in other models, we evaluated retinal function in eyes infected with WT or Δ*inhA1*, Δ*inhA2*, or Δ*inhA1-3 B. thuringiensis*. The absence of all three InhAs protected the eyes from retinal function loss, which would be expected in eyes with intact retinal structure and minimal inflammation (Figures 4, 8, and 12). Because WT and Δ*inhA1-3 B. thuringiensis* had similar cytotoxicity against RPE *in vitro*, the observed differences in retinal damage *in vivo* may not have been due to differences in the production of other toxins by these strains. Instead, these differences may correspond with the decreased bacterial growth and delayed inflammation in eyes infected with the strain lacking the InhA metalloproteases. As previously mentioned, the InhAs may have a role in growth via nutrient acquisition from the surrounding environment, which could explain the triple mutant’s delayed growth *in vivo*, and, subsequently, its attenuated retinal damage and inflammation.

This study is the first to address the importance of the *Bacillus* metalloproteases in an experimental intraocular eye infection, which mimics human infection. We demonstrated that a deficiency in *Bacillus* InhAs resulted in an attenuated intraocular infection. Our findings suggest that the InhAs may facilitate intraocular bacterial growth by producing an environment more conducive to persistence. Current therapeutics are relatively ineffective in treating *Bacillus* endophthalmitis [1,9–11,59–63], and due to its rapidly blinding course, *Bacillus* endophthalmitis requires early and precise treatment [62,63]. Treatment strategies for *Bacillus* and other types of endophthalmitis should be based on knowledge of the virulence factors that contribute to disease pathogenesis. Therefore, these metalloproteases may prove to be ideal targets for therapeutics in this potentially blinding disease.

## Materials and Methods

### Ethics Statement

The described experiments were conducted following the guidelines in the *Guide for the Care and Use of Laboratory Animals*, the Association for Research in Vision and Ophthalmology Statement for the Use of Animals in Ophthalmic and Vision Research, and the University of Oklahoma Health Sciences Center Institutional Animal Care and Use Committee (approved protocol 18-043).

### Bacterial Strains

*B. thuringiensis* 407 (WT) or its isogenic InhA mutants (Δ*inhA1*, Δ*inhA2*, Δ*inhA1-3*) [35,64] were injected into mouse eyes, as previously described [2,6,21,23,26,65,65–67]. Phenotypes of WT, Δ*inhA1*, Δ*inhA2*, and Δ*inhA1-3 B. thuringiensis* were compared by quantifying growth, hemolysis, cytotoxicity, and proteolysis, as described below.

### Bacterial Growth Curves

All strains were cultured for 10 hours with aeration at 37°C in brain heart infusion (BHI; VWR, Radnor, PA, USA) medium. Strains were diluted in fresh BHI to approximately 10^5^ CFU/mL and incubated for an additional 10 hours. Every 2 hours, 20-μL aliquots were track diluted in PBS and plated onto BHI agar plates, and counted after 24 hours [2,6,21,23,26,65,65–67]. Growth rates were analyzed by the equation, *N_t_* = *N*_0_ × (1 + *r*)^*t*^, where N_t_ is the concentration of bacteria at the end time, N_0_ is the concentration of bacteria at the initial time, r is the growth rate, and t is the time passed.

### Hemolytic Analysis

WT and Δ*inhA B. thuringiensis* were each cultured as described above, then centrifuged at 4150 rpm for 10 minutes. Supernatants were removed and filter sterilized (0.22 μm; Millex-GP, Merck Millipore Ltd., Cork, Ireland) and diluted two-fold in PBS (pH 7.4). Dilutions were incubated 1:1 with 4% (vol/vol) sheep red blood cells (Rockland Immunochemicals, Pottstown, PA, USA) in a round-bottom microtiter plate for 30 minutes at 37°C. After centrifugation at 300 x *g* for 10 minutes, supernatants were transferred into a flat-bottom microtiter plate and hemoglobin release was quantified (490 nm, FLUOstar Omega, BMG Labtech, Cary, NC, USA). Values are expressed in percent hemolysis relative to a 100% lysis control, as previously described [16,19,26,61,68].Values represent the means ± SD of two independent experiments.

### Retinal Pigment Epithelial Cell Cytotoxicity

Human ARPE-19 cells (American Type Culture Collection, Manassas, VA, USA) were grown to confluence in culture medium (DMEM/F12 supplemented with 10% fetal bovine serum and 1% glutamine; Gibco, Grand Island, NY, USA), diluted, and seeded into sterile 24-well plates. Twenty-thousand cells/100 μL were seeded in triplicate wells for overnight incubation. Supernatants of WT and Δ*inhA B. thuringiensis* were generated as described above, diluted 1:2, and added to wells containing ARPE-19. Cytotoxicity was measured by quantifying lactate dehydrogenase (LDH) from ARPE-19 (Pierce LDH Cytotoxicity Assay Kit, ThermoFisher Scientific, Waltham, MA, USA), according to the manufacturer’s instructions [12,20].

### Proteolytic Activity Assay

Protease production of WT and Δ*inhA B. thuringiensis* was detected on skim milk agar plates (CRITERION™; Hardy Diagnostics, Santa Maria, CA, USA). Colonies were transferred to the skim milk agar plates using a sterile toothpick and were incubated at 37°C for 48 h. Clear zones with at least 1 mm around single colonies indicated the activity of secreted proteases capable of hydrolyzing casein. Proteolytic zones were measured in millimeters from the edge of the colony to the edge of the clear zone.

### Mice and Intraocular Infections

*In vivo* experiments used mice (C57BL/6J, stock No. 000664; Jackson Labs, Bar Harbor, ME, USA). Mice were housed on a 12-hour light/dark cycle, for at least 2 weeks to equilibrate their microbiota, and under biosafety level 2 conditions. Mice (8-10 weeks old) were sedated with a cocktail of ketamine (85 mg/kg body weight; Ketathesia, Henry Schein Animal Health, Dublin, OH, USA) and xylazine (14 mg/kg body weight; AnaSed; Akorn Inc., Decatur, IL, USA). Deep anesthesia was confirmed by toe pinch. Infections were initiated by intravitreal injection containing ~200 CFU WT or Δ*inhA B. thuringiensis* in 0.5 μL BHI using a sterile glass capillary needle, as previously described [2,6,21,23,26,65,65–67]. Uninjected fellow eyes served as controls. As described below, eyes were analyzed by electroretinography (ERG) prior to euthanasia. After euthanasia (CO_2_ inhalation), eyes were harvested and infection courses were analyzed by quantifying intraocular *Bacillus* and polymorphonuclear leukocyte (PMN) infiltration (myeloperoxidase [MPO] activity), and histology.

### Quantifying Intraocular Bacterial Growth

Intraocular *Bacillus* were quantified from harvested eyes at 0, 2, 6, 8, 10, 12, 14, and 16 hours postinfection. Harvested eyes were homogenized in 400 μL PBS containing sterile glass beads (1 mm; BioSpec Products, Inc., Bartlesville, OK, USA). Eye homogenates were then track diluted onto BHI agar and counted, as previously described [2,6,21,23,26,65,65–67].

### Electroretinography

ERG was used to quantify retinal function in eyes infected with WT or Δ*inhA B. thuringiensis*, as previously described [2,6,21,23,26,65,65–67]. Scotopic ERGs were performed at 6, 8, 10, 12, 14, and 16 hours postinfection using Espion E2 software (Diagnosys LLC, Lowell, MA, USA). Mice were dark-adapted for at least 6 hours prior to ERG. Mice were anesthetized as noted above, and pupils were topically dilated (Phenylephrine HCl 2.5%; Akorn, Inc.). Two gold wire electrodes were placed on each cornea and reference electrodes were place on the forehead and tail. Eyes were then stimulated by five flashes of white light (1200 cd·s/m^2^). A- and B-wave amplitudes were recorded for both eyes in the same animal. The percentage of retinal function retained was calculated using the formula 100 – {[1 – (experimental A-wave amplitude/control A-wave amplitude)] x 100} or 100 – {[1 – (experimental B-wave amplitude/control B-wave amplitude)] x 100}.

### Histology

For histology, eyes were harvested from euthanized mice at 6, 8, 10, and 12 hours postinfection, incubated in low-alcoholic fixative for 30 minutes, and transferred to 70% ethanol for at least 24 hours. Paraffin-embedded eyes were sectioned and stained with hematoxylin and eosin [2,6,21,23,26,65,65–67].

### Estimation of Inflammatory Cell Influx

PMN infiltration into eyes was estimated by quantifying MPO using a sandwich ELISA (Hycult Biotech, Plymouth Meeting, PA, USA) as previously described [6,27,65,66]. Eyes were harvested at 6, 8, 10, and 12 hours postinfection, transferred into PBS-containing proteinase inhibitor (Roche Diagnostics, Indianapolis, IN, USA), and homogenized using sterile glass beads as described above. Eye homogenates were assayed with the MPO ELISA. The lower limit of detection for this assay was 2 ng/mL.

### RNA Isolation and Quantitative PCR

Expression of immune inhibitor A metalloprotease (InhA1, InhA2, and InhA3) genes from WT or Δ*inhA B. thuringiensis* was measured by real-time quantitative PCR (rtQPCR). Strains were grown in BHI. Total RNA was isolated from 18-hour cultures (RNeasy Mini Kit; QIAGEN, Hilden, Germany), DNA was removed (TURBO DNA-free Kit; Invitrogen, Carlsbad, CA, USA), and RNA was purified (RNA Clean & Concentrator-5 Kit; Zymo Research, Irvine, CA, USA), all following kit manufacturer’s instructions. RNA purity and concentration was confirmed using a Nanodrop (ThermoFisher). rtQPCR was performed (Applied BioSystems 7500; ThermoFisher), using the iTaq Universal SYBR Green One-Step Kit (Bio-Rad, Hercules, CA, USA) and primers listed in Table 1. Amplifications were performed in triplicate. Relative gene expression was determined using the ΔCT method, using 16S rRNA as a reference housekeeping gene.

**Table 1.**
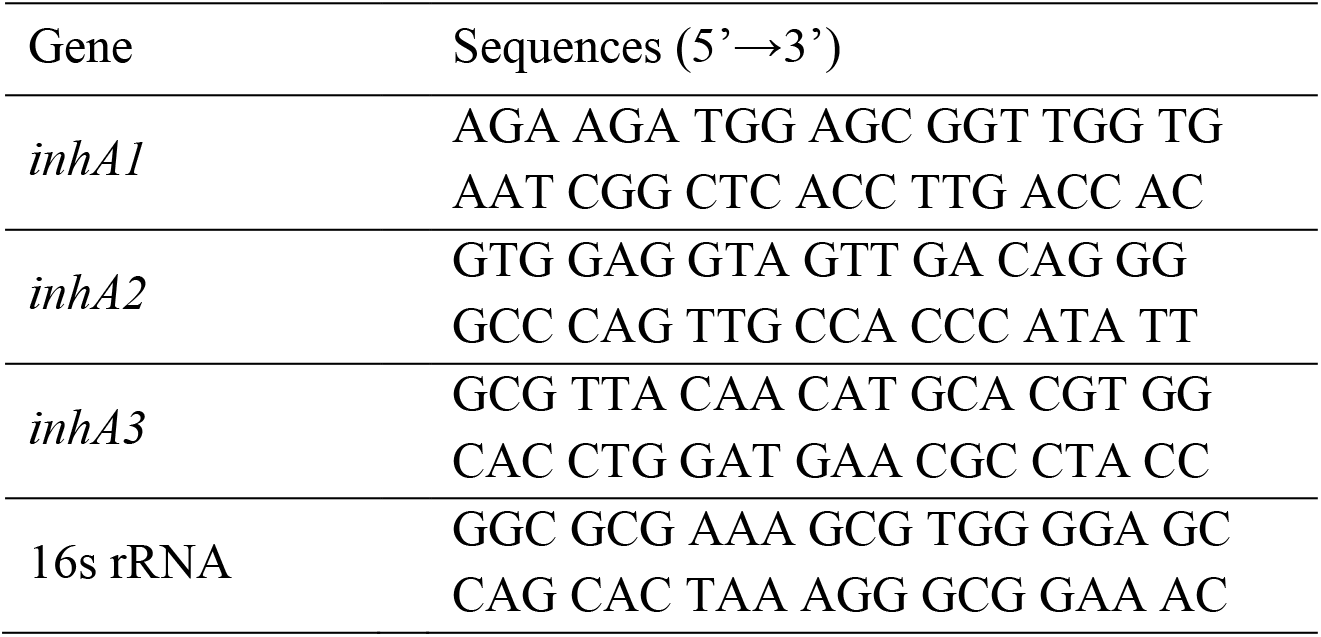
Primers Used in Quantitative PCR

### Statistics

Mann-Whitney U test was used for statistical comparisons unless otherwise specified (GraphPad Prism 7 Software, Inc., La Jolla, CA, USA) [26,66,68]. P values of < 0.05 were considered significant.

## Acknowledgements

The authors thank Dr. Feng Li and Mark Dittmar (OUHSC Vision P30 Live Animal Imaging and Analysis Core, Manali Kamath and Wil Nightengale for assistance with RNA preparations and myeloperoxidase experiments. Dean A. McGee Eye Institute, Oklahoma City, OK, USA) for invaluable technical assistance, and Excalibur Pathology (Moore, OK, USA) and the OUHSC P30 Cellular Imaging Core (Dean A. McGee Eye Institute, Oklahoma City, OK, USA) for histology expertise. This work was presented in part at the Association for Research in Vision and Ophthalmology Annual Meeting 2019, Vancouver BC and at the Society for Leukocyte Biology Annual Meeting 2019, Boston MA.

This work was supported by National Institutes of Health grants R01EY028810 and R01EY024140 (to MCC). Our research was also supported in part by National Institutes of Health grants R01EY025947 and R21EY028066 (to MCC), National Eye Institute Vision Core Grant P30EY021725 (to MCC), a Presbyterian Health Foundation Research Support Grant Award (to MCC and MHM), a Presbyterian Health Foundation Equipment Grant (to Robert E. Anderson, OUHSC), and an unrestricted grant to the Dean A. McGee Eye Institute from Research to Prevent Blindness.

